# Airway Basal Cells Show Regionally Distinct Potential to Undergo Metaplastic Differentiation

**DOI:** 10.1101/2022.05.04.490677

**Authors:** Yizhuo Zhou, Ying Yang, Jun Qian, Jian Ge, Debora Sinner, Hongxu Ding, Andrea Califano, Wellington V. Cardoso

**Author notes:** These authors contributed equally (Yizhuo Zhou, Ying Yang). **Author Contributions:** Y.Y and Y.Z. conceived the study. Y.Y performed single cell sequencing. Y.Z., Y.Y. and J.Q. performed immunofluorescence staining. Y.Z. performed organotypic and organoid cultures, q-PCR and all *in vivo* injury experiments. J.G performed polidocanol injury experiments. Y.Y. and Y.Z. performed quantification and statistics. HD performed computational analysis. W.V.C., Y.Y., H.D., A.C and Y.Z. interpreted the results, analyzed the data, produced the figures, and wrote the manuscripts. **Competing Interest Statement:** The authors declare no competing interests.

## Abstract

Basal cells are multipotent stem cells of a variety of organs, and in the lung are known as crucial components of the airway epithelium. However, it remains unclear how diverse basal cells are and whether distinct subpopulations respond differently to airway challenges. Using single cell RNA-sequencing and functional approaches, we report a significant and previously underappreciated degree of heterogeneity in the basal cell pool, leading to identification of six subpopulations in the murine trachea. Among these we found two major subpopulations comprising the most stem-like progenitor compartment, but with distinct signatures and ability to self-renew and differentiate. Notably, these occupy distinct ventral and dorsal tracheal niches and differ in their ability to initiate an aberrant program of differentiation in response to environmental perturbations in primary cultures and in injury mouse models *in vivo*. We found that such heterogeneity is acquired prenatally, when the basal cell pool and local niches are being established, and depends on the integrity of these niches, as supported by the altered basal cell phenotype of cartilage-deficient mouse mutants. Lastly, we show that key features that distinguish these progenitor subpopulations in murine airways are conserved in humans. Together, the data provide critical insights into the origin and impact of basal cell heterogeneity on the establishment of regionally distinct responses of the airway epithelium during injury-repair and in disease conditions.

## INTRODUCTION

Basal cells (BCs) are tissue-specific, adult, multipotent stem cells of various organs, including skin, esophagus, olfactory and the airway epithelia^1,2^. In the respiratory tract, BCs are found largely as a continuous layer of cells, at the base of the pseudostratified epithelium of extrapulmonary airways in mice and extending to small intrapulmonary conducting airways in humans^1,3^. A large body of studies, over decades, has shown that BCs are essential for the homeostatic maintenance and repair of the respiratory epithelium. Dysregulated airway BC activities, by environmental exposures and repeated injuries, are known to result in aberrant self-renewal and epithelial differentiation, often associated with chronic obstructive pulmonary diseases (COPD) and pre-neoplastic lesions that can lead to lung squamous cell carcinoma^1,4,5,6^.

The abundance and crucial role of these cells in the airway epithelium have, for many years, raised questions about whether BCs may represent a homogenous or a molecularly diverse pool of stem cells. Initial clues suggesting BC heterogeneity came from studies showing that they could differ in their ability to retain label in bromodeoxyuridine (BrdU) incorporation assays, and exhibit distinct clonogenic behavior due to different ability to activate *Krt5* promoter^7,8^. This hypothesis was further strengthened by the identification of a subpopulation of *Krt14*+ BCs, shown by lineage tracing to rapidly expand to generate additional BCs but not other more differentiated components of the damaged airway epithelium post-injury^9,10^. Long-term clonal analysis using a *Krt5-CreERT2* mouse line in combination with mathematical modeling, suggested that airway BCs comprise two subpopulations of multipotent stem cells, present in equal numbers, which differ in their commitment from a *bona fide* stem cell to an intermediate luminal progenitor not particularly associated with a specific cell niche^11^. Mouse models of lung injury-repair have shown that, immediately after severe injury of the airway epithelium and prior to regeneration of the luminal cells, two discrete BC subpopulations can be identified and distinguished by expression of determinants of cell fate commitment to the secretory or ciliated cell lineage^12^.

Single cell RNA sequencing (scRNA-Seq) analysis and computational modeling approaches are now providing new insights into BC diversity in the adult lung and are helping describe the differences in BC populations as the manifestation of discrete, co-existing cell states. One such study reports two BC states in the adult human lung based on the level of immaturity of these cells distinguished by levels of P63 and NPPC (natriuretic peptide C)^13^. Another study, also in human lungs, stratifies BCs into multipotent, proliferating, primed secretory, and activated subpopulations^14^. A third study, reporting human BCs as proliferating, differentiating, or quiescent, shows differential spatial enrichment of these BC subpopulations, with proliferating and differentiating BCs found only in large airways while quiescent BCs populate both large and small airways^15^.

In mice, a combination of scRNA-Seq profiling and lineage tracing assays have provided new insights into cellular hierarchies and BC trajectories but have not stratified BC subtypes in terms of their spatial distribution in the tracheal epithelium^16,17^. Using a sampling criterion based strictly on their spatial distribution, distinct BC phenotypes have been identified in tissues isolated from the dorsal and ventral mouse trachea^18^.

Taken together, these observations support the notion that the basal stem cell pool is diverse, while leaving key unresolved questions. What is the actual spectrum of BC diversity in a resting, unperturbed state of the adult murine trachea? Does BC diversity translate into a distinct ability to regenerate or undergo aberrant repair in injured airways? Is there any evidence of conservation in these features between mouse and human airways?

To address these questions, we have performed unbiased analysis of single cell RNA-Seq profiles from the *p63-CreERT2; R26-tdTomato* reporter mice, under homeostasis, and in different *in vitro* and *in vivo* models of injury-repair. Our studies show a previously underappreciated BC heterogeneity in the adult murine airway, with six molecularly distinct BC subtypes broadly organized into two major subpopulations, uniquely localized in distinct regional domains of expression. Functional analyses showed that these two subpopulations respond differently when animals are subjected to injury and maintain their phenotype even when isolated from their niche. Lastly, we show that these features are acquired during embryonic development and are conserved in BC from human airways.

## RESULTS

### scRNA-Seq reveals six distinct BC subpopulations in adult murine trachea under homeostasis

Mouse models of lung injury have provided major insights into how BCs contribute to regenerating the airway epithelium. Nevertheless, there is still limited information on how diverse the BC stem cell pool is in the unperturbed adult murine trachea. Although previous studies using scRNA-Seq approach have provided clear evidence of BC heterogeneity, methodological differences, including choice of mouse lines for BC isolation, ability to capture all subpopulations and choice of computational analysis did not allow clear insights into the extent to which steady-state BCs differ^16,17^.

To have a more precise idea about the composition of the BC pool, we specifically sorted these cells using a *p63-CreERT2; R26-tdTomato* reporter mouse line, known to label universally all BC and thus with the great chance to encompass all subtypes. This choice was supported by extensive evidence of *p63* expression in BCs throughout the body and the inability of these cells to form in the absence of *p63*^19, 20, 21, 22, 23^. Adult *p63-CreERT2; R26-tdTomato* mice were administered tamoxifen (240 μg/g body weight) and examined 72h later. Co-staining of P63+ cells with tdTomato in the tracheal epithelium confirmed efficient labeling of all BCs **(Figure 1A-B)**. KRT5 largely overlapped with P63 and tdTomato, although rare KRT5+ P63-tdTom-suprabasal cells could be identified as previously reported^24,25^ **(Figure 1B)**.

**Figure 1.**
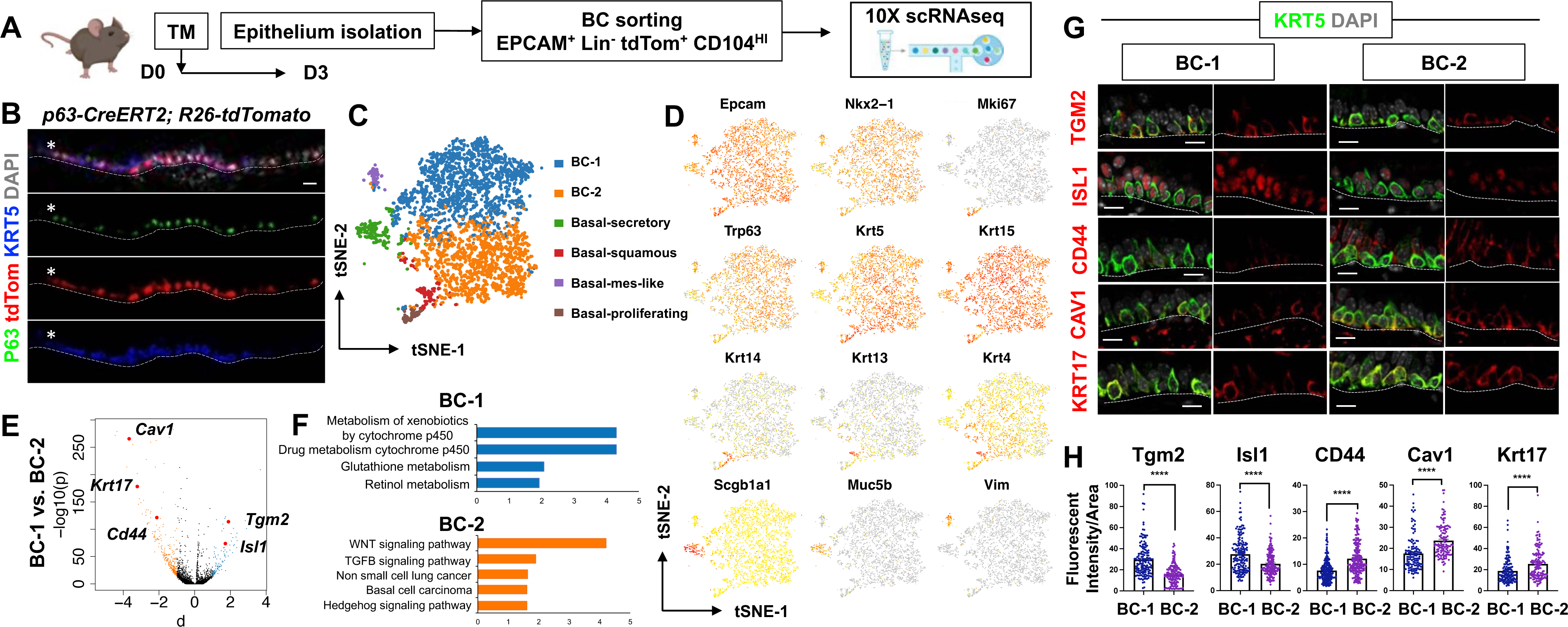
scRNA-Seq reveals mouse airway BC heterogeneity which can be mapped to distinct niches. (A) BC purifying strategy for scRNAseq using *p63-CreERT2; R26-tdTomato* lineage labeling and surface markers. (B) IF of tracheal sections showing efficient tdTom labeling in P63+ KRT5+ BCs. Asterisk denotes KRT5-P63+ suparabasal cell not labeled by tdTom. (C) tSNE visualization of airway BC scRNA-Seq, colored by cluster assignment and annotated using known lineage-specific markers. (D) tSNE visualization of airway BC scRNA-Seq, colored by expression (log_2_(TPM+1)) of each marker gene. **(E)** Volcano plot showing differentially expressed genes in Basal-1 and Basal-2 clusters represented in black dots; genes significantly enriched in Basal-1 were colored in blue (-log_10_(p) > 10; d (average expression difference) >1); genes significantly enriched in adult Basal-2 were colored in orange (-log_10_(q) > 10; d < -1). (F) Enriched KEGG gene sets in each subpopulation in Basal-1 (blue) and Basal-2 (orange). (G) IF of tracheal serial sections showing expression patterns of 5 identified markers co-labeling with KRT5. (H) Quantification of fluorescent intensities for 5 markers comparing two BC subpopulations residing in different tissue locations. n = 3 animals. Each dot represents a KRT5+ cells analyzed. 8 to 15 views for each location from 5 to 9 sections were imaged. Statistics: Student’s t-test, ****p < 0.0001. Scale bars represent 10 μm.

Thus, we sorted tdTomato^+^ EPCAM^+^ Lin^-^ CD104^HI^ airway BCs, validated their purity using cytospin **(Figure S1A-B)**, and generated scRNA-Seq profiles using the 10X Genomics Chromium. A total of 4,209 single cells passed quality control, with median values of 10,802 transcripts and 2,956 genes per cell detected (see Methods). Using the Cellranger toolkit (10X Genomics), followed by marker gene annotation, we identified six distinct BC subpopulations, which were confirmed to originate from the respiratory lineage (*Epcam*+ *Nkx2-1*+) and to express established adult BC markers (*p63, Krt5, Aqp5, Aqp3*, and *Pdpn*) (**Figure S1C**).

Cluster analysis showed that these subpopulations differed widely in their contribution to the BC pool. Notably, two clusters were prominently represented, accounting for 85.8% of sequenced BCs. They were found at an approximate 1:1 ratio and were named as BC-1 (46.1%) and BC-2 (39.7%). Both lacked *Krt14* expression, while expressing comparable levels of canonical BC markers (*p63, Krt5, Krt15*). Luminal lineage markers were nearly absent in these clusters and secretory genes (*Scgb1a1, Scgb3a1, Scgb3a2)* were detected at very low levels **(Figure 1C-D, Figure S1-2)**. Based on their signatures, BC-1 and BC-2 appeared to be the most stem-like BC subpopulations and did not seem to overlap with any of the BC subpopulations reported by previous scRNA-Seq studies. **(Figure 1C-D, Figure S1-3)**.

The remaining four clusters comprised only a small fraction of the total BC pool (14.2%). Based on their gene signature, featuring markers associated with differentiated cell phenotypes, these were classified as Basal-proliferating (*Mki67, Ccna2, Aurkb, Ube2c*: 2.1%), Basal-secretory (*Scgb1a1, Scgb3a1, Scgb3a2, Muc5b*: 5.8%), Basal-squamous (*Krt13, Krt4, Krt6a, Sprr2a3*: 3.3%), and a mixed cluster, as represented by expression of genes associated with ciliated, neuroendocrine and mesenchymal cells, which together accounted for 3.1% of all profiled cells **(Figure 1C-D, Figure S1-2)**.

### Two major BC subpopulations reside in distinct niches

Since BC-1 and BC-2 constituted the majority of the BCs in the airway epithelium, we further examined their transcriptional signatures to assess their differences, including against other BC subpopulations. Despite some similarities, when compared to other BC subpopulations, BC-1 and BC-2 presented a markedly distinct gene expression signature **(Figure S1-S2, S4)**. Gene set enrichment analysis of KEGG pathways revealed enrichment of BC-1 cells in drug metabolism, glutathione and retinol metabolism signatures. By contrast, BC-2 were differentially enriched in Wnt, TGF beta and Hedgehog signaling pathways signatures, as well as in basal cell carcinoma and non-small cell lung cancer^26,27,28,29^.

We identified the most differentially expressed genes between BC-1 and BC-2 (-Log10(p) ≥ 10; d (average expression difference) > 1) and selected representative candidates for further characterization of their sites of expression in the adult trachea by immunofluorescence (IF) assays at the encoded protein level. Candidates were selected based on statistical significance and validated antibody availability. Based on these criteria we mapped the expression of *Tgm2* (transglutaminase-2, C polypeptide), *Isl1* (ISL LIM homeobox), *Cav1* (caveolin1), *Cd44* (Cd44 antigen, or Pgp1) and *Krt17* (keratin-17, K17) in tracheal sections of adult mouse lungs **(Figure 1E)**.

Co-IF with a KRT5 antibody confirmed BC-specific expression of each of these makers in the upper and lower trachea with no clear evidence of anterior-posterior differential distribution of BC subpopulations. However, BC-1-specific markers *Tgm2* and *Isl1* were strongly expressed in ventral epithelium BCs and only weakly detected in the dorsal epithelium. By contrast, BCs of the dorsal trachea were marked by strong expression of *Cd44, Cav1, Krt17*, which were only weakly expression in ventral BCs (**Figure 1G**). Quantitative analysis of fluorescence intensity confirmed the significant difference in signals for these markers between the BC subpopulations (**Figure 1H**).

### BC-1 and BC-2 are functionally different and maintain their distinct ability to self-renew *in vitro*

To further investigate the phenotypic differences between BC-1 and BC-2 cells, we asked whether their unique gene signatures were dependent on active signals from their respective local microenvironments in the trachea. Given the major differences in tissue structure and gene networks between the dorsal and ventral tracheal mesenchyme, we examined the extent to which BC-1 and BC-2 preserved their identities when isolated and expanded in culture. For this, whole tracheas from adult mice were dissected, and the membranous (dorsal) and cartilaginous (ventral) regions were surgically separated and processed for isolation and analysis of BC cultures from each sample. Approximately equal amounts of cells were seeded in Transwell plates and expanded under submerged conditions, as previously described^25^.

No obvious differences in growth were observed between BCs from the two regions as they expanded, and both reached confluency by day 0 in culture (**Figure 2A**). Double-IF of the above-described BC-1 and BC-2 markers with KRT5 showed a signal in cultures from both groups. However, TGM2 and ISL1 IF was clearly stronger in ventral BC cultures, while CAV1, CD44 and KRT17 IF was distinctly stronger in dorsal BC cultures (**Figure 2B**). qRT-PCR analysis was consistent with the *in vitro* and *in vivo* IF patterns (**Figure 2C**). These observations support the hypothesis that, *in vitro*, BC-1 and BC-2 cells isolated from the adult trachea can maintain their phenotype independent of their respective microenvironment for at least a week, thus providing an opportunity to assess their functional differences as adult progenitors of the airway epithelium.

**Figure 2.**
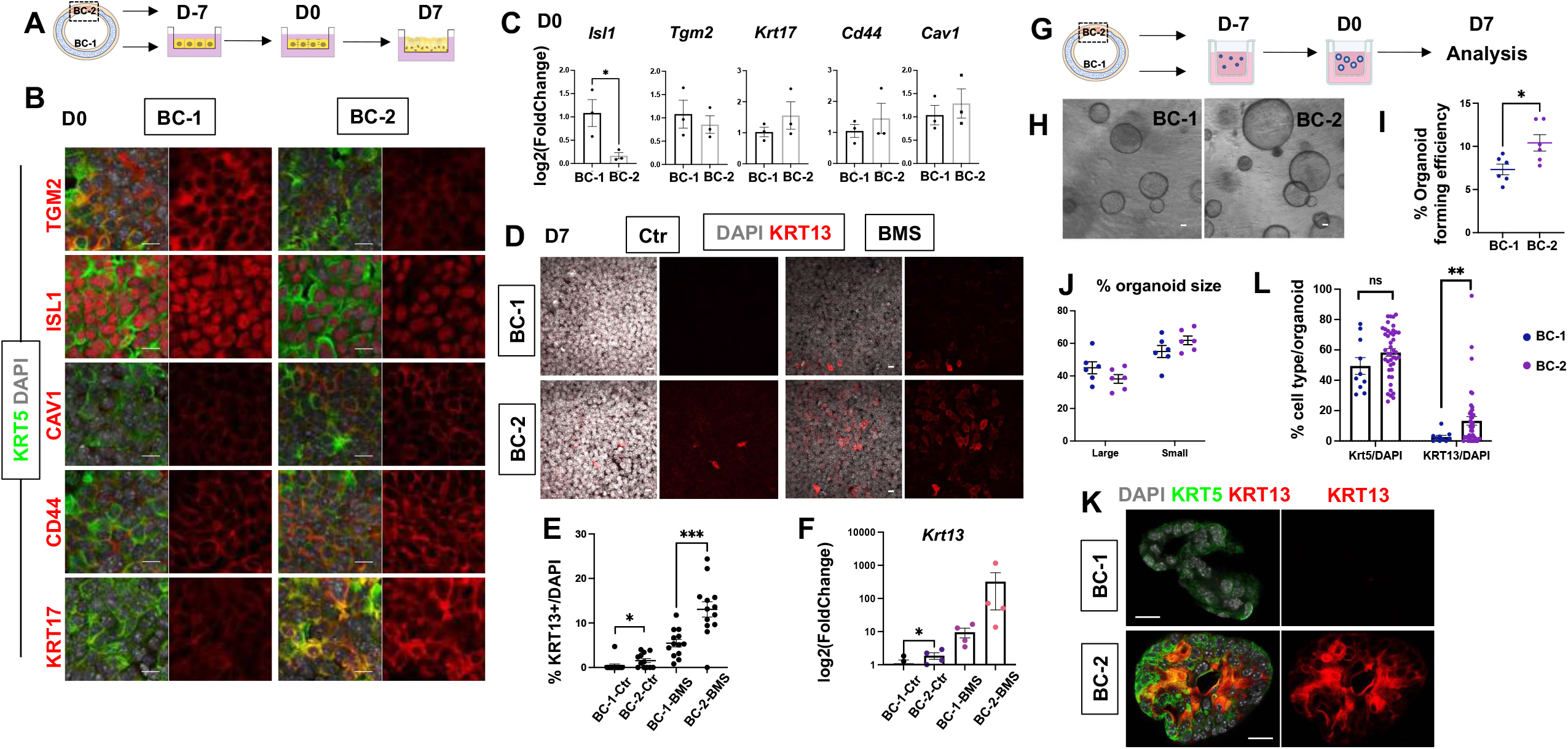
BC-2 are more prone to squamous differentiation in ALI and organoid culture. (A) Schematic representation about isolation of BCs from distinct tissue locations for ALI culture. (B) IF of 5 new markers in KRT5+ BCs at D0 after 7-day submerged culture. (C) qPCR of 5 new markers for D0 culture derived from BC-1 or BC-2. Data are normalized to BC-1. Bar graph representing mean±SEM, dots representing each replicate. n = 3. (D) IF of KRT13 for D7 ALI culture with or without BMS treatment. (E) Morphometric analysis of KRT13+ cells for D7 ALI culture with or without BMS treatment. Scattered dots representing %KRT13+ cells in total population from each field, with mean±SEM from 4 fields per batch, n = 3 batches for each condition. (F) qPCR of *Krt13* for D7 ALI culture with or without BMS treatment. Data are normalized to BC-1 control (without BMS treatment). Bar graph representing mean±SEM, dots representing each replicate. n = 4 batches. (G) Schematic representation about isolation of BC-1 and BC-2 for organoid culture. (H) Bright view of BC-1 and BC-2 derived organoids at D7. (I) Scattered dots representing % organoid number in seeding cell number, with mean±SEM from 6 fields for each condition. (J) Scattered dots representing % small/large-sized organoids in BC-1 or BC-2 derived culture, with mean±SEM from 6 batches for each condition. See methods for quantification details. (K) IF of organoid section showing abundant KRT13 in mBC-derived organoids. (L) Morphometric analysis of % KRT5+ and KRT13+ cells in each organoid. Bar graph representing mean±SEM, dots representing each organoid, BC-1: n = 10. BC-2: n = 43. Statistics: Student’s t-test, *p < 0.05, ***p < 0.001. Scale bars represent 10 μm.

To identify differences in progenitor cell behavior, BC-1 and BC-2 cells from ventral and dorsal trachea were isolated (as above), embedded in 3D Matrigel cultures in Transwell plates, and allowed to form tracheospheres (**Figure 2G**)^3^. Analysis of these cultures on D7 confirmed the ability of both populations to form organoids, albeit at different efficiency. Quantitative analysis revealed a significantly higher number of tracheospheres derived from the dorsal (BC-2) cultures compared to their ventral (BC-1) counterpart (**Figure 2G-I**, 10.41±0.009% versus 7.32±0.006%, respectively). Next, we examined whether BC-1 and BC-2-derived tracheospheres differed in size. Morphometric analysis was performed in D7 cultures from both groups (n = 6 batches per condition) and organoids were classified according to size into large (≥ 35.65 μm) and small (< 35.65 μm) (see methods and figure legend). We found no statistically significant differences in the average size of large and small organoids from BC-1 and BC-2 cultures (**Figure 2J)**.

Taken together, these data suggest that the observed differences in transcriptional signature between the two main BC subpopulations play a functional role, since BC-2 appear better equipped to initiate a program of expansion to form a higher number of tracheospheres.

### BC-1 and BC-2 differ in their ability to initiate an aberrant program of differentiation in primary cultures

To identify additional functional features that may distinguish these two progenitor cell subpopulations. we studied genes—especially transcription factors—that were most differentially expressed between BC-1 and BC-2 cells. The analysis revealed significant enrichment in genes associated with squamous cell differentiation. BC-mediated generation of a squamous stratified epithelium is a key event in the normal differentiation trajectory of the skin, esophagus and other organs. In the lung, however, it is the hallmark of an aberrant program triggered by sustained epithelial injury giving rise to squamous metaplasia, a reversible event that can however also lead to pre-malignant lesions^30,31,32^. Among the transcription factors differentially expressed and upregulated in BC-2 we found the Krüppel-like Factors *Klf4, Klf6* and *Klf13, Barx2*, AP1 family members (*JunB, JunD, ATF3*), reported to promote squamous cell commitment in stratified epithelia^30,33,34,35,36,37,38,39^. BC-2 also differed from the BC-1 cluster as a result of its enrichment in in squamous differentiation genes from the Hedgehog/Tgfb/Wnt pathways **(Figure 1E-F, Figure S3A-B**). Importantly, GSEA revealed enrichment of a smoking-related squamous metaplasia signature in BC-2, but not in BC-1 cells^40^ **(Figure S4C)**.

The cluster analysis had already identified a small subpopulation of Basal-Squamous cells featuring expression of a *Krt13, Krt4, Krt6a*, and *Sprr2a3* marker signature, which is characteristic of differentiated squamous cells (**Figure 1C-D, Figure S1C, S2, S5**). Intriguingly, none of these genes could be found in the 100 most differentially-expressed in BC-2 compared to BC-1 (**Figure S4A**), despite BC-2 enrichment in regulators of the squamous cell differentiation program. More surprisingly, several genes whose expression was previously reported in BCs isolated from the dorsal trachea of adult mice, such as *Krt6a, Serpinb2* and *Krt13*^18^, were found in our BC-Squamous cluster (**Figure S5A-C**) but were not differentially expressed in BC-2 cells, which localize to the dorsal trachea. Taken together, these data suggest that, under homeostasis, adult tracheal BCs consist mostly of two subpopulations of multipotent progenitors, which remain largely uncommitted yet present distinct differentiation potential.

We speculated that BC-2 was more sensitive to environmental perturbations and more prone to respond by activating an aberrant program of differentiation. To explore this possibility, tracheospheres were generated with BCs isolated either from ventral or dorsal regions (largely representative of BC-1 and BC-2 subpopulations, **Figure 2G-L**) as before. After day 0, culture conditions were changed by shifting from a serum-based (MTEC Plus) to a serum-free (MTEC/SF) medium and the culture period was extended for an additional week. We then examined whether the prolonged time/SF conditions served as a perturbation sufficient to highlight differences in the differentiation potential BC-1or BC-2-derived organoids. Cultures were interrupted at day 7 and analyzed for their ability to maintain their KRT5+ progenitor cell pool and to induce KRT13, a marker of noncornified squamous epithelium in multiple organs, also known to label initiation of a metaplastic program in the airway epithelium of mice, rats and humans^41^. *Krt13* was also chosen as it featured as a topmost upregulated gene in the Basal-Squamous cluster and thus a key representative indicator of a squamous cell fate choice.

IF staining and quantitative analysis of KRT5 in day 7 organoids showed no difference in the relative number of labeled cells between groups (BC-1: 49.43±5.54%, BC-2: 58.29±2.50%, p = 0.17), suggesting similar self-renewal capacity (**Figure 2K-L**). By contrast, the number of KRT13+ cells was markedly different (**Figure 2K-L**. BC-1: 2.65±1.08%, BC-2: 13.26±2.86%, p = 0.001), suggesting that, under similar environmental perturbation (changes in culture conditions), BC-2 may activate an aberrant differentiation program. This differential behavior was also observed when BC-1 and BC-2 were similarly expanded in MTEC Plus media and then allowed to differentiate as 2D ALI cultures in serum-free (MTEC/SF) medium. ALI day 7 BC-2 cultures expressed higher levels of *Krt13* expression and higher percentage of Krt13+ cells compared to BC-1-derived cultures (**Figure 2D-F**. 1.54±0.47% and 0.39±0.39% in BC-2 and BC-1, respectively).

In the respiratory epithelium squamous metaplasia is strongly linked to perturbations in the progenitor cell program triggered by disruption in the retinoic acid (RA) signaling, which has been extensively reported as a manifestation of Vitamin-A deficiency in humans and in animal models, as well as in cells cultured in the presence of pharmacological antagonists of the RA pathway^42,43,44,45^. Given the differential expression of retinoid-related genes between BC-2 and BC-1 cells, we examined how these progenitors responded when cultured in the presence of the pan-retinoic acid receptor inhibitor BMS-493 (BMS). IF staining of ALI day 7 cultures showed marked increase in the number of KRT13+ cells (BMS, 5.46±0.85% for BC-1 versus 13.06±1.72% for BC-2. **Figure 2D-E)**, as well as a trend towards upregulation of this gene and other squamous markers in BMS-treated BC-2-derived cultures compared to BC-1 cultures **(Figure 2F, Figure S6**)

Thus, scRNA-Seq analysis identified two major subpopulations of BCs in the adult trachea that occupied distinct niches along the dosal-ventral axis and present distinct susceptibility to activating an aberrant metaplastic program of differentiation under similar environmental perturbation *in vitro*.

### BC-2 shows markedly distinct behavior during initiation of repair of the damaged epithelium in mouse models of injury

These intriguing observations led us to inquire whether severe injury of the airway epithelium in the intact animal may reveal intrinsic differences in BC-1 vs. BC-2-mediated repair programs. For this, we used two classical mouse models of lung injury-repair *in vivo*. Intraperitoneal injection of Naphthalene (275 mg/kg body weight), a byproduct of the cigarette smoke metabolized by cytochrome P_450_, results in massive injury and sloughing of club cells of the airway epithelium. In the trachea, repair is subsequently initiated by activation of a BC program of regeneration that repopulates the epithelium within 2-3 weeks^46^. Polidocanol is a detergent/sclerosing agent that induces extensive epithelial sloughing when injected intratracheally in mice, leading to BC-mediated regenerative response to reconstitute the airway epithelium^47^.

Injury was thus induced with Naphthalene or Polidocanol in 8 week-old adult mice and animals were euthanized at different stages of the repair process for each injury model **(Figure 3A**). IF was performed for KRT5 and SCGB1A1 as representative markers of the basal and the regenerating luminal (secretory) compartment, and KRT13 was used to identify differences in the BC program, suggestive of a metaplastic state **(Figure 3)**. IF staining was performed in tracheal sections oriented for simultaneous exposure of both the ventral and dorsal tracheal epithelium, based on their association with cartilage or a smooth muscle layer.

**Figure 3.**
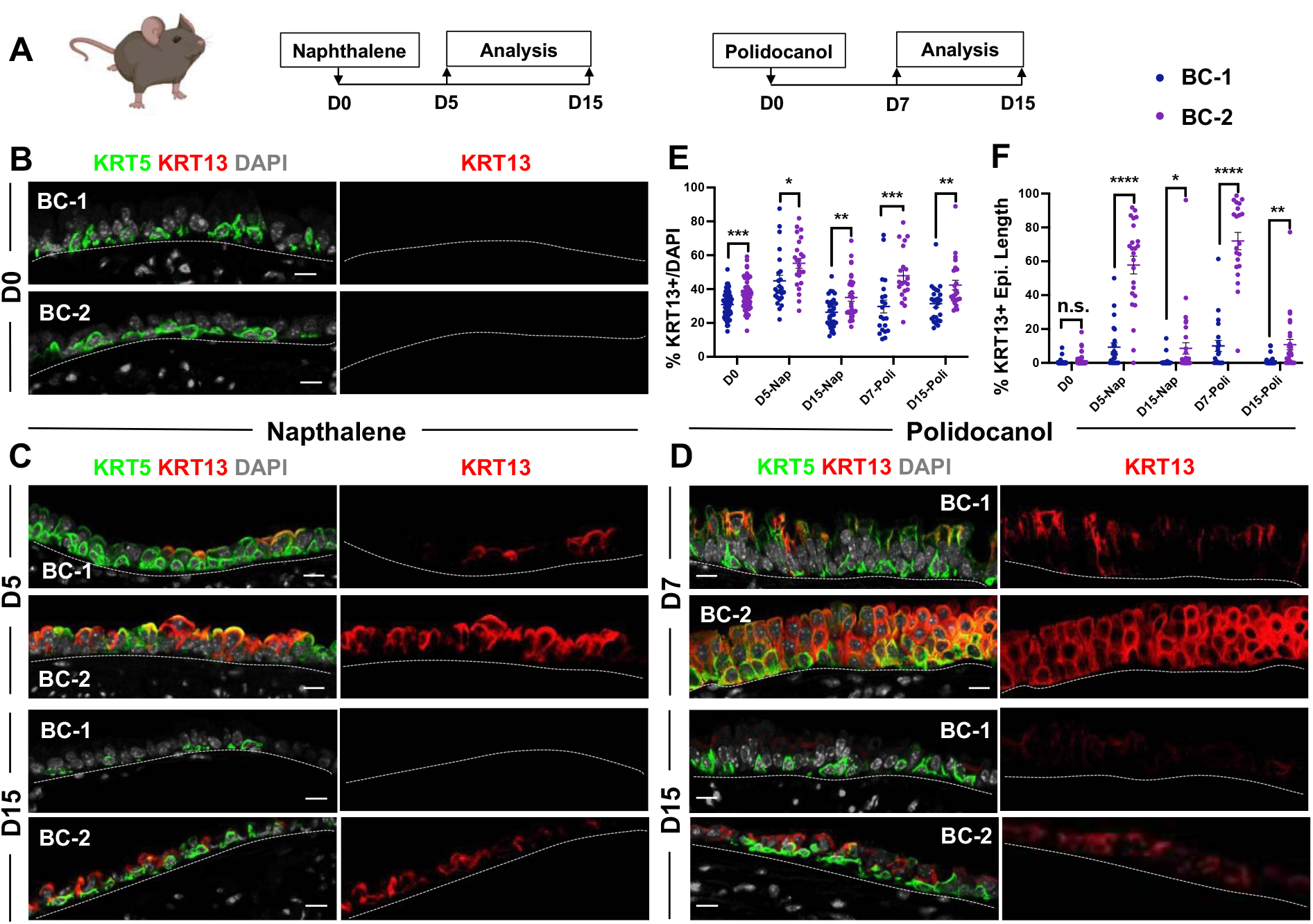
BC-1 and BC-2 behave differently in mouse airway injury-induced squamous metaplasia. (A) Naphthalene/Polidocanol-induced airway injury analyzed at early (D5/7) and late (D15) stages. (B-D) IF of tracheal sections at D0 as control (n = 4 animals), D5/7 (n= 3 animals) and D15 (n = 3 animals) post injury showing BC canonical marker KRT5 and squamous marker KRT13. (E) Morphometric analysis of KRT13+ cells for cartilage side or muscle side at D0, D5/7 and D15. Scattered dots representing % KRT13+ cells in total population from each field, with mean±SEM from n >= 22 paired views of different tissue locations. (F) Scattered dots representing % length of epithelium containing KRT13+ cells in total epithelium from each field, with mean±SEM from n >= 21 paired views of different tissue locations. Statistics: Student’s t-test, *p < 0.05, **p < 0.01, ***p < 0.001, ****p < 0.0001; n.s., not significant. Scale bars represent 10 μm.

In uninjured (control) animals, KRT13+ cells were only rarely observed in the tracheal epithelium, but increased significantly as epithelial regeneration initiated in both Naphthalene and Polidocanol injury models. Notably, differences in KRT13 labelling between the dorsal and ventral epithelium were strikingly accentuated as BCs initiated epithelial repair. At day 5 post-Naphthalene the number of KRT13+ cells increased dorsally from 0.71±0.37% (control) to 45.47±4.74% of all epithelial, while ventrally they increased from 0.2±0.12% (control) to 8.36±2.55%. Similarly, at day 7 post-polidocanol injury KRT13+ comprised 53.41±5.33% of all the dorsal regenerating epithelium, compared to only 8.69±2.46% ventrally (**Figure 3B, 3C-D upper panel, 3E**). Consistent with this observation, quantitative analysis of the perimeter occupied by KRT13+ cells along the basement membrane in the regenerating epithelium was significantly higher dorsally, compared to the ventral trachea (Naphthalene: 57.79±5.28% vs. 9.25±2.64%; Polidocanol 71.98±5.13% vs. 10.03±3.28%, **Figure 3F**). Interestingly, regenerating club cells— as identified by SCGB1A1 expression—were only infrequently detected in the dorsal epithelium, where KRT13+ cells were abundant. In these areas KRT13 was not only co-labeled with KRT5 in the basal layer, but also found widely expressed in the nascent suprabasal/luminal cells of the dorsal epithelium. This contrasted with the strong SCGB1A1 signals in the KRT13 low epithelium indicating robust induction of club cell regeneration in the ventral epithelium at this stage (**Figure S7)**. Thus, KRT13 and SCGB1A1 emerged as inversely correlated in the BC-derived regenerative program at this stage.

Surprisingly, by day 15 post injury, when the epithelium had already extensively regenerated in both models, the proportion of KRT13+ labeled cells was significantly reduced in both the dorsal and ventral compartments (Naphthalene: 5.79±2.19% vs. 0.44±0.34%; Polidocanol:10.69±2.53% vs. 0.54±0.31%, **Figure 3B, 3C-D lower panel, 3E)**. The areas contiguously occupied by KRT13 expressing cells along the basement membrane were also markedly reduced by day 15 in the dorsal and ventral trachea (Naphthalene: 8.56±3.35% vs. 0.72±0.5%; Polidocanol: 10.8±3.01% vs. 1.19±0.54%, **Figure 3F)**.

Overall, these data suggest that the distinct BC subpopulations identified by an unbiased scRNA-Seq analysis could be identified not only by their transcriptional signature, but also by their spatial distribution, ability to form organoids, and induction of *Krt13*, a gene associated with a metaplastic program in airway BCs. This KRT13-associated program appeared to be transiently induced and reversible in the absence of persistent stimulus.

### BC heterogeneity is defined during embryonic development

A key observation from our analyses was that the identity and intrinsic behavior of BC-1 and BC-2 cells were maintained in culture, independent of their niche, for at least 14 days. We thus asked whether acquisition of heterogeneity was an early event that paralleled BC fate specification during lung development. We have previously shown that, in the embryonic mouse lung, airway epithelial cells destined to become BCs (pre-basal cells) are already specified at an early stage, when airways are still forming, but have to undergo a series of maturation events to become properly functional ^48^. Thus, to investigate the developmental basis of the BC heterogeneity, we first searched for evidence of differential distribution of the BC-1 and BC-2 markers in Pre-BCs from ventral and dorsal embryonic tracheas. Tracheas were isolated from E18.5 embryos and triple-labeled for KRT5 with each marker and a-SMA (as a reference for the dorsal location). This revealed significant differential enrichment of BC-1 and BC-2 markers, consistent with the pattern identified in the adult trachea. Quantitative analysis of IF intensity, specifically in KRT5-expressing cells, confirmed statistically significant difference of expression for all markers except CAV1 (See methods, **Figure 4A, 4C)**.

**Figure 4.**
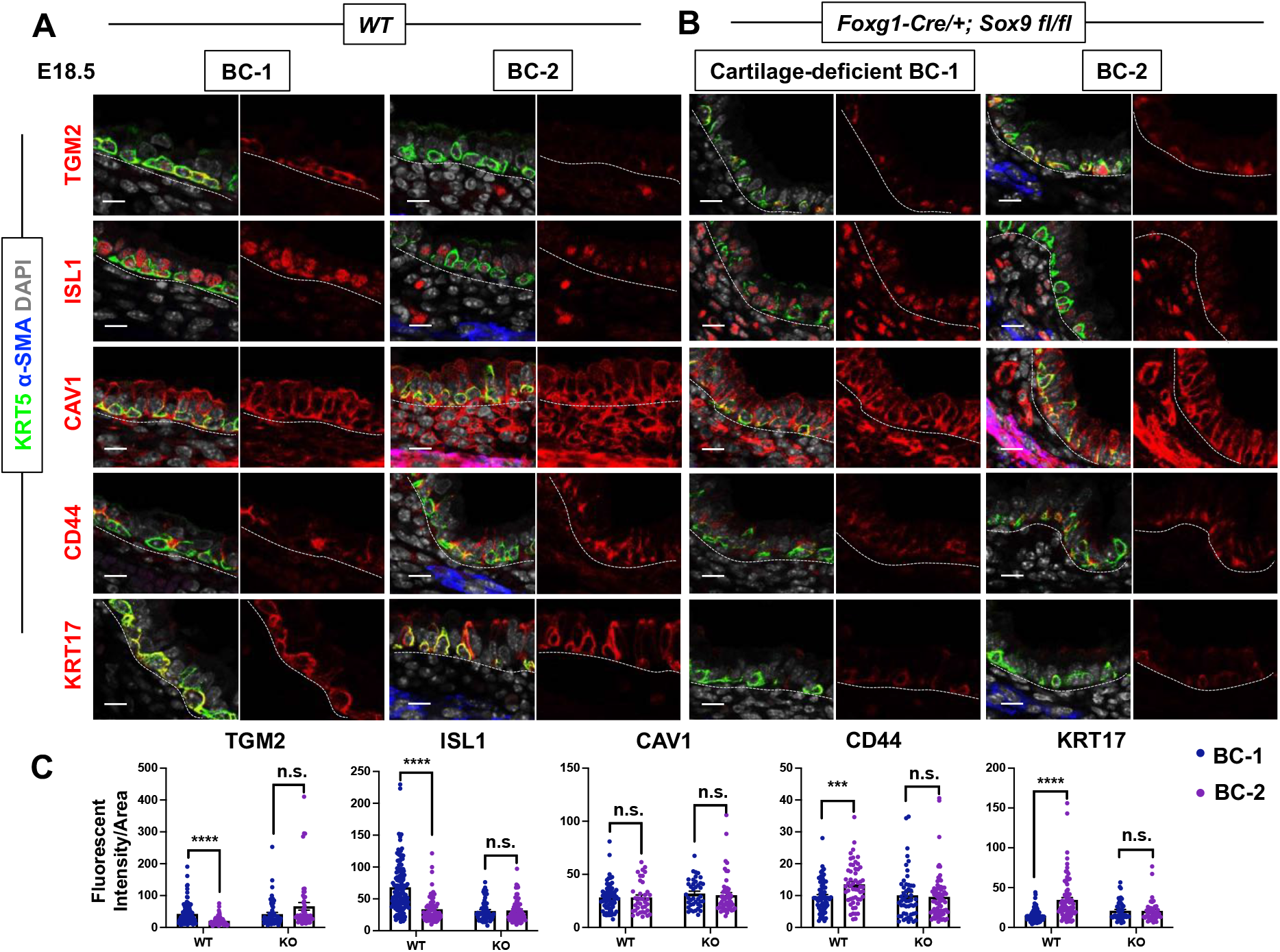
BC-1/2 spatial heterogeneity is shaped by the niches during development. (A-B) IF of tracheal sections at E18.5 in *WT* littermate control and *Foxg1-Cre/+; Sox9 fl/fl*. 5 new markers are co-stained with canonical BC marker KRT5 and smooth muscle marker a-SMA. (C) Fluorescent intensity quantification for the 5 new markers on different tissue locations. Bar graph representing mean±SEM, dots representing each cell, n = 3 animals for each genotype. Statistics: Student’s t-test, ***p < 0.001, ****p < 0.0001; n.s., not significant. Scale bars represent 10 μm.

Next, we reasoned that local epithelial-mesenchymal crosstalks were crucial in establishing BC heterogeneity, as they were being specified and underwent maturation in the developing trachea. Cartilage primordia appear early in tracheal development at around E10.5 when basal cell precursors are already recognizable but have not yet acquired *Krt5* expression and undergone lineage restriction^48,49^. Genetic studies have demonstrated the crucial role of *Sox9* as a master regulator of cartilage cell fate^50^. Targeted deletion of *Sox9* in the developing tracheal mesenchyme of *Foxg1-Cre; Sox9 fl/fl* mutants results in complete absence of cartilage, a tracheal defect lethal at birth^49,51,52,53^.

The inability of these mutants to specify cartilage cell fate and to establish a ventral niche while BCs were developing provided a unique opportunity to examine the impact of the local environment in BC-1 vs BC-2 diversification. Tracheas were isolated from WT and *Foxg1-Cre; Sox9 fl/fl* mutants at E18.5 and analyzed for the dorsal-ventral distribution of the BC-1/BC-2 markers by IF, as described above. We restricted our analysis to E18.5 because of the low levels or inconsistent expression of these markers at earlier stages. As previously reported, tracheal rings or cartilage condensations were absent in *Foxg1-Cre; Sox9 fl/fl* mutants and replaced by an indistinct mesenchymal tissue in the ventral trachea, while, dorsally, a smooth muscle layer was evident^49,51^. IF staining for KRT5 identified Pre-BCs in both ventral and dorsal regions of these mutants, confirming previous reports that disruption of the cartilage niche may lead to an overall decreased number of Pre-BCs (**Figure 4A-B**). Tracheal sections triple-labeled with each marker/KRT5/a-SMA showed signals in both ventral and dorsal pre-BC of E18.5 WT and mutants. Remarkably, a comparison of the intensity of signals between ventral and dorsal Pre-BCs from mutants showed no significant difference for all markers, in sharp contrast with the findings in WT tracheas (**Figure 4A-C**). This appears to have resulted from an attenuation of signals in the ventral pre-BCs from the mutants, suggesting that cartilage-derived signals may be key in creating the differences in gene expression between BC-1 and BC-2.

The data underscored a key role for epithelial-mesenchymal crosstalks during co-development of the cartilage and the BC pool, generating spatial heterogeneity among pre-BCs of the embryonic trachea.

### Airway BC spatial heterogeneity is conserved in human airways

scRNA-Seq analysis of human lung BCs has shown them to comprise a heterogenous population of multipotent progenitors. However, subtype numbers, as well as criteria for grouping and localization vary across studies. We asked whether the markers of BC-1 and BC-2 subpopulations identified in the murine trachea were also differentially enriched in spatially distinct BC subpopulations of the human airways.

IF staining of the five representative markers of BC-1 and BC-2 in sections of healthy adult human lung donors showed BC expression of *ISL1, CAV1, CD44, KRT17* but not *TGM2*. The latter was rather present in some luminal cells (**Figure 5A**). Quantitation of fluorescent intensity of these signals in BCs showed a remarkably conserved pattern of differential enrichment with higher levels of ISL1 in ventral BCs, contrasting with the higher levels of CAV1, CD44, and KRT17 in dorsal BCs (**Figure 5A-B**). Notably and consistent with our findings in mice, these features continued to distinguish the two BC populations even when they were isolated from their original niche and maintained in culture under the same conditions. IF staining of human BCs expanded to confluency showed distinctly stronger ISL1 signals in the cultures isolated from ventral (BC-1) in contrast to the dorsal BC-2-derived cultures which were readily recognizable by the stronger CAV1, CD44 and KRT17 expression **(Figure 5C-E)**.

**Figure 5.**
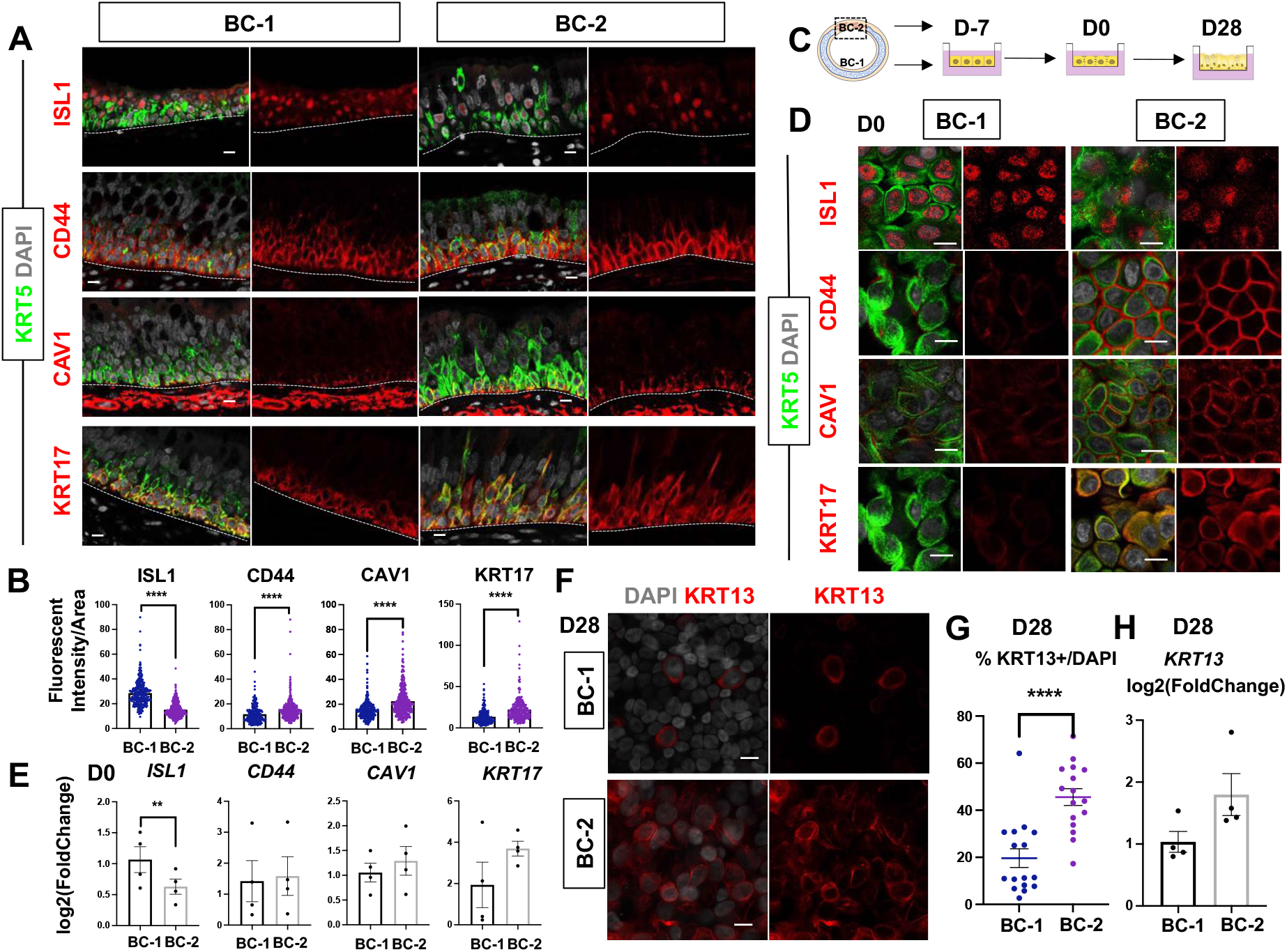
BC-1/2 spatial heterogeneity is conserved in human. (A) IF of human large airway sections for ISL1, CD44, CAV1 and KRT17 co-stained with KRT5. (B) Fluorescent intensity quantification for ISL1, CD44, CAV1 and KRT17 on paired different tissue locations. Bar graph representing mean±SEM, dots representing each cell, n >= 227 from 3 different human donors. (C) Schematic representation about isolation of BC-1 and BC-2 for ALI culture. (D) IF of 4 markers in KRT5+ BCs at D0 after 7-day submerged culture. (E) qPCR of 4 markers for D0 culture derived from BC-1 or BC-2. Data are normalized to BC-1. Bar graph representing mean±SEM, dots representing each replicate. n = 3 from one donor. (F) IF of KRT13 for D28 ALI culture. (G) Morphometric analysis of KRT13+ cells for D28 ALI culture. Scattered dots representing %KRT13+ cells in total population from each field, with mean±SEM from 5-6 fields per batch, n = 3 batches from one donor. (F) qPCR of *KRT13* for D28 ALI culture. Data are normalized to BC-1. Bar graph representing mean±SEM, dots representing each replicate. n = 4 batches from one donor. Statistics: Student’s t-test, **p < 0.01, ****p < 0.0001; n.s., not significant. Scale bars represent 10 μm.

Furthermore, when allowed to differentiate under ALI human dorsal BC-derived cultures consistently generated more KRT13+ cells (**Figure 5F-H**, 19.67±3.98% compared to 14.31±3.58%). Differences between cultures from these distinct BC subpopulations could be seen even after prolonged time, as shown in ALI day 28 cells. These observations support the idea of potentially conserved mechanisms of BC diversification between mice and humans. A better understanding of these issues could provide information crucial for the development of models relevant to human lung repair-regeneration.

## Discussion

In the present study, we report a broader range of heterogeneity of BCs in the progenitor cell pool of the adult murine airways under homeostasis. We show that in spite of this diversity, most of the BCs comprise two major subpopulations which are similarly uncommitted, compared to the other BC subpopulations. These uncommitted progenitors exhibit comparable levels of canonical BC markers but have distinct signatures and differ from each other in their ability to initiate a metaplastic program of differentiation. Remarkably, their intrinsic differences in phenotype appear to be largely determined by their spatial localization and established when BCs are still developing in the embryonic trachea. Once established, airway BCs can maintain their subtype-specific features to adulthood and behave distinctly in response to different perturbations *in vitro* as well as injury challenges to airway epithelium *in vivo*.

As adult stem cells, airway BCs continuously self-renew and differentiate into luminal lineages during homeostasis. Rather than characterizing BC diversity based on their differentiation potential predicted by trajectory analysis, here we identified the various subpopulations based on their bona fide signature in an unperturbed state. The use of *p63-CreERT2; R26-tdTomato* mice in our scRNA-Seq screen allowed the identification of a broader range of BCs compared to similar screens using *Krt5-CreER* based lines, given previous reports of KRT5 negative cells in the BC pool^48,54^. Recent scRNA-Seq studies identify a panel of BC markers that includes *Hlf, Icam1, Notch1, Ngfr, Krt15, F3, Aqp3*, besides *Krt5* also present in our screen^17,16^. While we found them widely distributed in all clusters none of these distinguished the two main BC subpopulations identified in our study (**Figure S3**).

Spatial heterogeneity has been reported in human BCs isolated from nasal, trachea and intrapulmonary airways with region-specific phenotypes shown to be maintained in culture over extended time^55^. A surprising finding from our report was the unbiased identification of these two main BC subpopulations differentially distributed in the dorsal and ventral trachea in agreement with the findings from a recent study in which BCs were isolated and analyzed primarily from these regions^18^. Our data support the idea that heterogeneity is assigned while BCs are specified and may involve epigenetic modifications. Detailed characterization of BC-1 and BC-2 at homeostasis on chromatin accessibility, histone modification, DNA methylation may further unveil the molecular regulatory mechanisms of BC heterogeneity. scRNA-Seq of the corresponding niches will also provide valuable information about intercellular signal communications critical for BC heterogeneity

Phenotypic differences have been reported in BCs of the tracheal epithelium associated with cartilage rings compared to those present in the intercartilaginous regions in which the mesenchyme exhibits a smooth muscle layer^56,57,58^. While both BCs are present in the ventral epithelium, our markers could not consistently distinguish cartilage-associated from the intercartilage BC populations. This could be ascribed to the fact that the BC-1 and BC-2 markers found from our unbiased approach were expressed in both ventral and dorsal subpopulations at different levels, but not enough to consistently distinguish cartilaginous vs non-cartilaginous BCs in our assays. The functional assays in organoid and organotypic cultures were also not designed to identify these differences since in these preparations both subpopulations of ventral BCs were mixed.

The potential clinical significance of these finds is intriguing as they point to a differential susceptibility of the BC-1 and BC-2-derived epithelia to activate a potentially pathological program associated with *Krt13*. Squamous metaplasia is a common pre-neoplastic lesion in smokers and patients with chronic obstructive pulmonary diseases (COPD). Although reversible, under continued injury the metaplastic epithelium can undergo malignant transformation to lung squamous cell carcinoma. Interestingly, several genes found to be differentially enriched in the BC-2 subpopulation, such as *Tppp3, Tnfrsf12a*, and *Cav1* have been associated with severity or aggressiveness of squamous carcinoma in various organs^59,60,61^. Future studies exploring the impact of BC heterogeneity in the molecular mechanisms of initiation of tumorigenesis can provide significant insights into the pathogenesis of human conditions.

## Acknowledgements

We would like to thank all members of Cardoso lab and the members of the CCHD, Dr. Jianwen Que, Dr. Munemasa Mori and Dr. Hans Snoeck for thoughtful discussions. We also thank Dr. Erin Bush from Columbia Genome Center for helping single cell sequencing. This work was supported by NIH -NHLBI R35-HL135834-01 to W.V.C., NIH-NHLBI RO1 144744 to D.S., and NIH S10OD020056 to CCTI (Columbia Center for Translational Immunology) Flow Cytometry Core and NIH/NCI Cancer Center Support Grant P30CA013696 to HICCC (Herbert Irving Comprehensive Cancer Center), and the National Center for Advancing Translational Sciences NIH-UL1TR001873 to CTSA (Clinical and Translational Science Award).

## METHODS

### Mice and tamoxifen induction of lineage labeling

*p63-CreERT2; R26-tdTomato* mice were generated and characterized as described in Lee et al. Int. J. Biol. Sci. 2014 and Yang et al. Dev Cell 2018. To label airway BCs at homeostasis, 6-12 week-old mice were exposed to 240 μg/g body weight TM once via oral gavage. A short chase period of 3 days was used to ensure accumulation of tdTomato in BCs and to minimize tdTomato labeling in luminal descendants.

To generate *Foxg1-Cre; Sox9 fl/fl* embryos, *Sox9 fl/fl* mice (Kist R. et al. Genesis, 2002)) were mated with *Foxg1-Cre/+* mice (Hebert JM, et al. Dev Biol. 2000) and subsequently rebred to generate *Foxg1-Cre; Sox9 fl/fl*; embryos as previously described in Nasr T. et al. Dis Model Mech, 2021 and Bottasso et al Am J Physiol Lung Cell Mol Physiol. 2022. Genotypes of transgenic mice were determined by PCR using genomic DNA isolated from mouse-tails or embryonic tissue (*Sox9* F: 5’ *CCG GCT GCT GGG AAA GTA TAT G* 3’. *Sox9* R: 5’ *CGC TGG TAT TCA GGG AGG TAC A* 3’. *Foxg1-Cre* F: 5’ *TGC CAC GAC CAA GTG ACA GCA ATG* 3’. *Foxg1-Cre* R: 5’ *AGA GAC GGA AAT CCA TCG CTC G* 3’). For embryonic developmental staging, the morning when vaginal plug was detected was considered E 0.5. Animals were housed in a pathogen-free environment and handled according to the protocols approved by CCHMC Institutional Animal Care and Use Committee (Cincinnati, OH, USA).

Wild-type C57/Bl6 6-12 week-old mice were used for naphthalene and polidocanol injury models. Only females were used for naphthalene injury. Both male and females were used for polidocanol injury. Details were described below.

All studies were approved by Columbia University Institutional Animal Care and Use committees (IACUC).

### Epithelial cell isolation and fluorescence activated cell sorting (FACS)

Tracheas and esophagus were isolated from adult mice with brief cleanup of stromal tissues. To isolate tracheal epithelial cells, only the main tracheal region below cricoid cartilage and above bifurcation was used to avoid confounding p63^+^ cells from larynx or mainstem bronchi. Tracheal tubes were cut open and digested in Dispase Digestion Solution (16U/ml Dispase (Corning, 354235) + 10μg/ml DNase I diluted in PBS) at room temperature for 40min. Digestion was stopped by transferring tracheas to PBS. Epithelium was physically peeled off with forceps, and collected in Falcon tubes. Further digestion into single cells was done in Trypsin Digestion Solution (0.1% Trypsin + 3mM EDTA diluted in PBS) at 37°C for 10min. To isolate esophageal epithelial cells, the muscle layer surrounding esophageal epithelium was physically peeled off, following with Trypsin digestion of epithelium at 37°C for 10min. Digestion was stopped by addition of FACS buffer (HBSS + 2% BSA + 1x GlutaMax) with 10% FBS, followed by gentle pipetting and passage through 70μm cell strainer.

To stain for cell sorting, cells were suspended in staining buffer (FACS buffer + 10μg/ml DNase I), and incubated with antibodies for 40min at 4°C. Cells were washed with FACS buffer and DAPI was added to a final concentration of 1.25μg/ml before sorting. Sorting was performed on Influx (BD Biosciences) and data analyzed with Flowjo (version 10). Cells were collected in staining buffer. The following antibodies were used: CD104-BV510 (1:50, BD Biosciences, 743079), EPCAM-APC (1:100, eBioscience, 17-5791-82), Lineage cocktail-Alexa Fluor 700 (1:20, Biolegend, 133313).

### Single cell RNA sequencing

Immediately after sorting, cells were stored on ice and processed at the Columbia University Genome Center for 10X Chromium preparation. Single Cell 3’ libraries were prepared using the Chromium Single Cell 3’ v2 Protocol (CG00052) according to the manufacturer’s manual (10X Genomics). The pooled, 3’-end libraries were sequenced using Illumina HiSeq4000. Cell Ranger version 2.1.1 was used for primary data analysis, including demultiplexing, alignment, mapping, gene expression quantification, dimension reduction analysis and clustering analysis within individual datasets. Specifically, for alignment and mapping, the mm10 reference genome and corresponding annotation were used.

### Gene set overlapping analysis

Basal-1/2 specific differentially expressed genes highlighted in Figure 1F (blue and orange, respectively) were used for the analysis. The analysis was performed using Broad Institute web-based gene set overlapping analysis tool, as avaliable at http://software.broadinstitute.org/gsea/msigdb/annotate.jsp. KEGG gene sets were included in the analysis, and FDR q-value threshold was set as 0.05.

### GSEA analysis

The squamous metaplasia signature gene set was generated by converting human smoking-associated squamous metaplasia genes (Goldfarbmuren, Nat Commun. 2020) to mouse homologs. Basal-1/2 gene expression values from these two clusters from scRNA-seq profiles were used for comparison. GSEA was performed following the developer’s protocol (GSEA version 4.1.0, available at: http://www.gsea-msigdb.org/gsea/index.jsp).

### Histology and immunofluorescent staining

Embryonic and adult tracheas were fixed in 4% paraformaldehyde in PBS at 4°C overnight. After washing in PBS, samples were processed for frozen or paraffin-embedding.

Immunofluorescence (IF) was performed in tissue sections (6-8mm) blocked with 10% horse serum and 0.3% TritonX-100 (Sigma) for 1 hour at room temperature (rt). Primary antibodies were incubated in 1% bovine serum albumin (Sigma) and 0.3% TritonX-100 at 4°C overnight or 2 hours at room temperature. Sections were then washed with PBS and incubated with Alexa Fluor-conjugated secondary antibodies (1:500) and NucBlue Live Cell ReadyProbes Reagent (DAPI) (Life Technology) for 1 hr. After washing, samples were mounted with ProLong Gold antifade reagent (Life Technology). When necessary, antigen unmasking was done using Citric Based solution (Vector Labs H-3300) heated in microwave. Mouse primary antibody staining was done using M.O.M kit (Vector Labs BMK-2202).

The following primary antibodies were used: rabbit anti-p63a (1:100, CST, 13109s); chicken anti-Krt5 (1:300, Biolegend, 905901); rabbit anti-Krt17 (1:10000, Abcam, ab53707); rabbit anti-Cav1 (1:100, Cell Signaling Technology, 3267S); rat anti-Cd44 (1:100, BD Biosciences, BD553131); rabbit anti-Tgm2 (1:50, Cell Signaling Technology, 3557S); rabbit anti-Isl1 (1:50, Abcam, ab109517).

The following secondary antibodies were used: donkey anti-rabbit (conjugated with Alexa Fluor 488, 568, 647); donkey anti-chicken (conjugated with Alexa Fluor 488); goat anti-chicken (conjugated with Alexa Fluor 488, 647); donkey anti-mouse (conjugated with Alexa Fluor 488, 568, 647); donkey anti-rat (conjugated with Alexa Fluor 488, 647). All secondary antibodies were purchased from Thermo Fisher Scientific or Jackson ImmunoReseach.

Confocal microscopy was performed using a Zeiss LSM 710 confocal microscope through 20X, 40X or 63X lens.

### Fluorescent Intensity Profile Analysis

To compare the fluorescent intensities of each marker in tracheal BCs located on ventral side (lined by cartilage rings) to those on dorsal side (lined with the smooth muscle layer), paired images with the same z axis were acquired in the same section using exactly same digital setting. For each set of paired views, 10 to 20 BCs were randomly picked, circled around Krt5 labeled cellular shape. Fluorescent intensity and area (mm2) for each cell was measured by Zen 2.3 lite software. The intensity value of each marker channel per unit area for each BC was calculated and analyzed for statistics. For each marker, 8 to 15 views from 5 to 9 sections from 3 different animals were analyzed

To compare the fluorescent intensities of each marker in the mutant (*Foxg1-Cre; Sox9 fl/fl*) and control mice at developmental E18.5, cross-sectional immunofluorescence staining images were acquired using confocal microscopy. Only BCs above cartilage on the ventral side and BCs above the smooth muscle on the dorsal side were picked for analysis in control mice. Only BCs above the smooth muscle on the dorsal side and BCs facing the smooth muscle on the ventral side were picked for analysis in mutant mice. The average fluorescence intensity was measured as described above.

### Trachea Basal cell isolation for ALI culture

Mouse and human adult tracheal basal cells were dissected in Pronase from muscle side or cartilage side respectively. For mouse ALI culture, BCs from different tissue location were submerge cultured in Tracheal Epithelial Cell (MTEC) Plus medium for 7 days for expansion (D-7 to D0) and in MTEC serum-free media for ALI culture for 7 days (D0 to D7). For human ALI culture, BCs from different tissue location were submerge cultured in Bronchial epithelial growth medium (BEGM) for 7 days for expansion (D-7 to D0) and in BEGM-based ALI medium for ALI culture for 28 days (D0 to D28).

### Mouse 3D organoid culture and quantification

Mouse tracheal basal cells were resuspended in MTEC/Plus medium mixed 1:1 with growth factor– reduced Matrigel, and seeded into Transwell insert. MTEC/Plus was added to the lower chamber and changed every other day. On day 7, MTEC/SF was placed in the lower chamber and changed every other day (Rock et al. 2009). The spheres per insert were counted on days 12 and fixed on day14.

To quantify organoid colony forming efficiency, the total number of organoids per well was counted 14 days after seeding (Rock et al. 2009). All the experiments were done by using EVOS m5000 microscope (Invitrogen) at 4X magnification. To quantify organoid size, diameter of all organoids in a random field per well was measured 14 days after seeding by using Fuji ImageJ (Gao et. al, 2015). To quantify the percentages of Krt13-positive and Krt5-positive cells per organoid, organoids were randomly picked by using confocal microscopy and analyzed by Zen 2.3 lite software.

### Q-PCR

RNA was extracted using QIAGEN RNeasy Mini Kit and cDNA was synthesized using the SuperScript IV First-Strand synthesis system (Thermo Fisher). Gene expression was assessed using TaqMan Fast Universal PCR Master Mix (Applied Biosystems) and analyzed on a Step-One Plus instrument (Applied Biosystems). At least 3 biological repeats were analyzed for each group.

### Naphthalene/Polidocanol injury model

For Naphthalene injury, single dose of naphthalene (dissolved in sunflower oil) was administered to female wild-type mice by IP injection at 275 mg/kg body weight to induce trachea injury. Freshly prepared naphthalene was administered before noon (Aleksandra et al., 2018). For Polidocanol injury, wild-type mice were anesthetized and received 20ul 2% polidocanol (freshly prepared in PBS) by oropharyngeal aspiration delivery following previously published protocols (Paul et al., 2014).

Animals were sacrificed 5/7/15 days after injury. Paired images from both dorsal and ventral sides with the same z-axis were acquired in the same section using the same digital setting to quantify the percentages of Krt13-positive and Krt5-positive cells.

### Quantification and statistical analyses

Quantification and statistical analysis have already been detailed in the Methods section above, associated with each experiment, as well as in the figure legends. All quantification for colocalization and marker analyses were performed in Adobe Photoshop. Statistical analyses were performed in Microsoft Excel or GraphPad Prism.

## Code availability

Custom scripts for reproducing the figures are available upon request to the corresponding authors.

## Data availability

scRNA-Seq data for mouse trachea BCs described in the manuscript have been deposited at the Gene Expression Omnibus (GEO) under accession number GSE134064. IF staining images described in the manuscript were available upon request to the corresponding authors.

