## Supplementary Figures for "Airway Basal Cells Show Regionally Distinct Potential to Undergo Metaplastic Differentiation"

**Figure S1, related to Figure 1.** (A) Gating strategy to sort BCs for scRNA-Seq. (B) IF of cytospin of sorted airway BCs, stained with canonical markers KRT5 and P63. (C) Expression levels (log2(TPM+1)) of cluster-distinct genes (rows) in each epithelial cell (columns).

**Figure S2, related to Figure 1.** (A) Violin plots of scRNA-Seq clusters for general epithelial markers *Epcam*, *Nkx2-1*, canonical BC markers *Trp63*, *Krt5*, *Krt15*, squamous markers *Krt14*, *Krt13*, *Krt4*, secretory markers *Scgb1a1*, *Scgb3a2*, *Muc5b*, mesenchymal markers *Vim*, *Thy1*, cell cycle markers *Mki67*, *Aurkb*. (B) Bar plots of scRNA-Seq clusters for the per single cell Basal/Luminal scores (integrated z-scores, computed using Stouffer's method on per-gene z-scores) based on manually curated gene lists (Basal: *Trp63, Krt5, Snai2, Egfr, Krt15, Jag2, Dll1, Ngfr, Dapl1, Gpr87, Dlk2, Bcam*; Luminal: *Krt8, Scgb1a1, Scgb3a2, Muc1, Krt13, Krt4, Lypd2, Notch2, Notch3, Perp, Ly6d, Ly6e, Lypd3, Hes1*).

**Figure S3, related to Figure 1.** tSNE visualization of airway BC scRNA-Seq for BC markers identified from previous publications, colored by expression ($log_2$(TPM+1)) of each marker gene.

**Figure S4, related to Figure 1. BC subtypes showing unique gene expression patterns.** (A) Top 50 genes significantly enriched in BC-1 and BC-2 showing markedly difference in expression pattern in these two clusters. **(B)** BC-1 and BC-2 specific transcription factors. Relative expression levels of genes (row-wise Z-score of mean $log_2$(TPM+1)). (C) GSEA showing that BC-2 cluster exhibit strong correlation with squamous metaplasia signature.

**Figure S5, related to Figure 1. Squamous cluster showing unique gene expression patterns comparing to the rest cells.** (A) Volcano plot showing differentially expressed genes in Squamous cluster and the rest cells represented in black dots; known squamous markers are marked in red. (B) Enriched KEGG gene sets in genes significantly increased in the Squamous cluster (red) and significantly decreased in the Squamous cluster (blue). (C) Squamous cluster gene signature. Top 50 genes ranked by fold change significantly enriched in Squamous cluster comparing to the rest cells. (D) TFs specifically upregulated and down regulated in Squamous cluster comparing to the rest cells. Relative expression levels of genes (row-wise Z-score of mean log2(TPM+1)).

**Figure S6, related to Figure 2.** qPCR of additional squamous markers for D7 ALI culture with or without BMS treatment. Data are normalized to BC-1 control (without BMS treatment). Bar graph representing mean±SEM, dots representing each replicate. n = 4.

**Figure S7, related Figure 4.** (A) Naphthalene/Polidocanol-induced airway injury analyzed at early (D5/7) and late (D15) stages. (B-D) IF of tracheal sections at D0 as control, D5/7 and D15 post injury showing BC canonical marker KRT5 and secretory marker CC10.

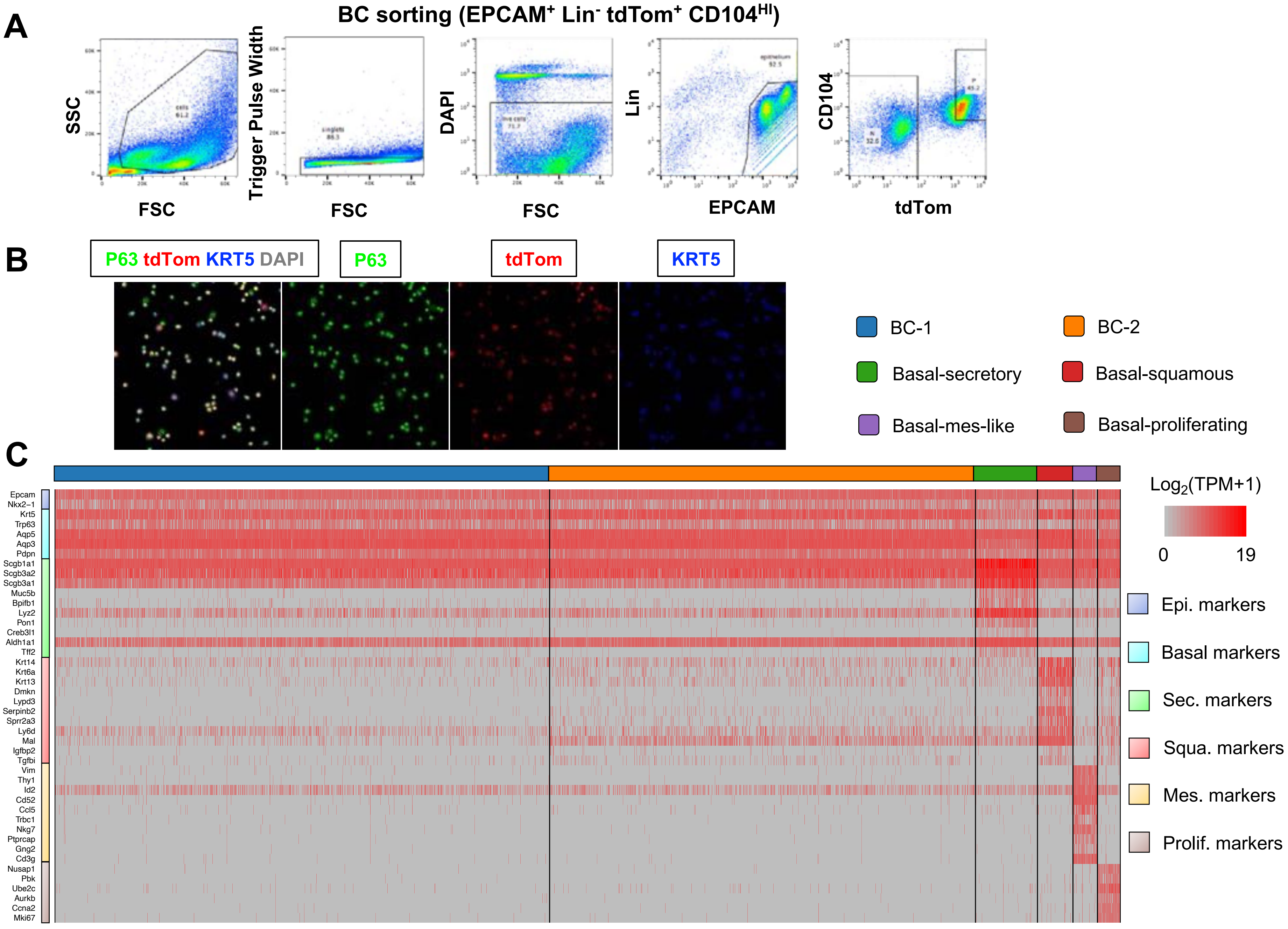
A
BC sorting (EPCAM+ Lin- tdTom+ CD104HI)
SSC
FSC
Trigger Pulse Width
FSC
DAPI
FSC
Lin
EPCAM
CD104
tdTom
B
P63 tdTom KRT5 DAPI
P63
tdTom
KRT5
BC-1
BC-2
Basal-secretory
Basal-squamous
Basal-mes-like
Basal-proliferating
C
Log2(TPM+1)
0
19
Epi. markers
Basal markers
Sec. markers
Squa. markers
Mes. markers
Prolif. markers
Epcam
Nkx2−1
Krt5
Trp63
Aqp5
Aqp3
Pdpn
Scgb1a1
Scgb3a2
Scgb3a1
Muc5b
Bpifb1
Lyz2
Pon1
Creb3l1
Aldh1a1
Tff2
Krt14
Krt6a
Krt13
Dmkn
Lypd3
Serpinb2
Sprr2a3
Ly6d
Mal
Igfbp2
Tgfbi
Vim
Thy1
Id2
Cd52
Ccl5
Trbc1
Nkg7
Ptprcap
Gng2
Cd3g
Nusap1
Pbk
Ube2c
Aurkb
Ccna2
Mki67

**Figure S1**

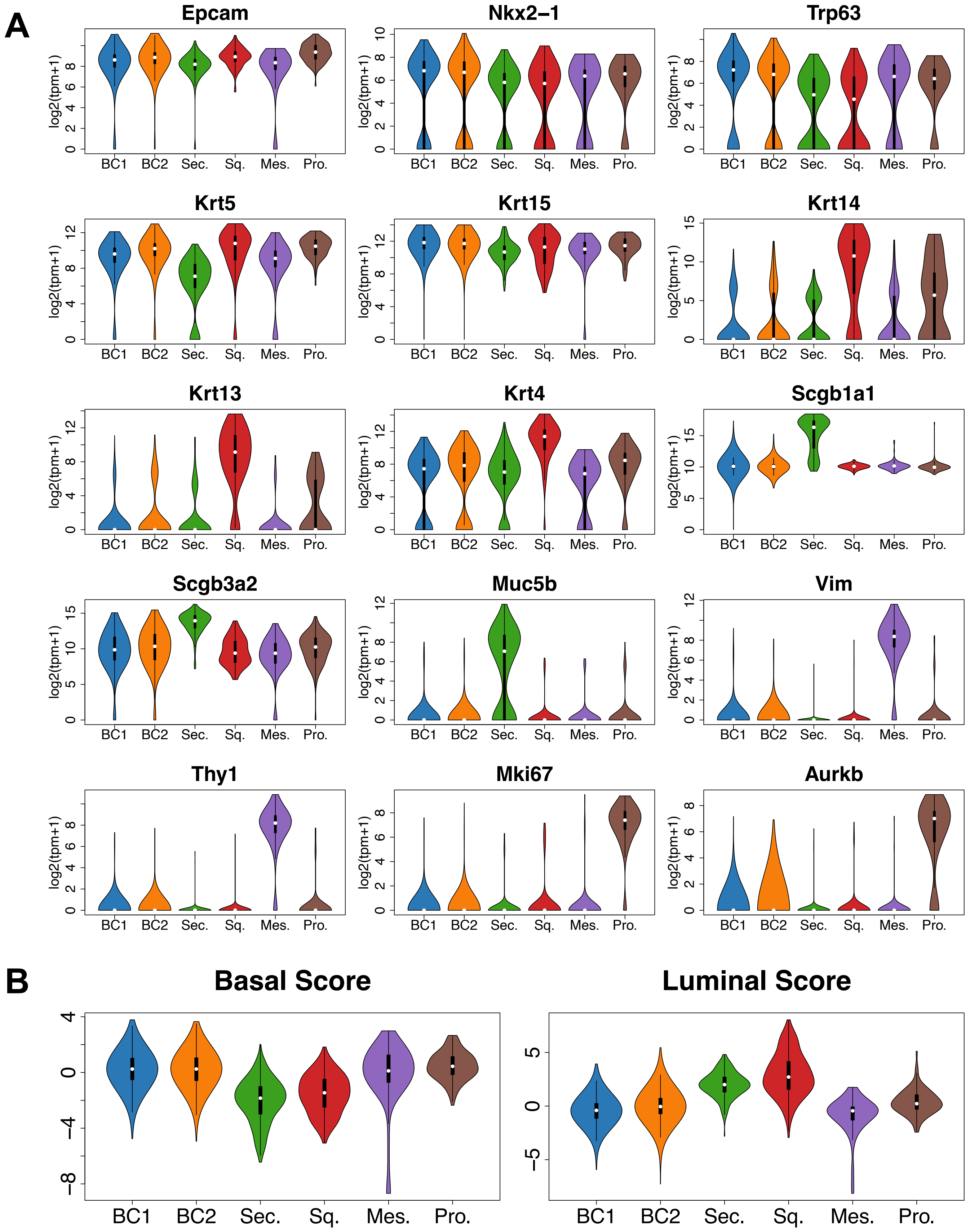
A
Epcam
Nkx2–1
Trp63
Krt5
Krt15
Krt14
Krt13
Krt4
Scgb1a1
Scgb3a2
Muc5b
Vim
Thy1
Mki67
Aurkb
log2(tpm+1)
BC1
BC2
Sec.
Sq.
Mes.
Pro.
B
Basal Score
Luminal Score

**Figure S2**

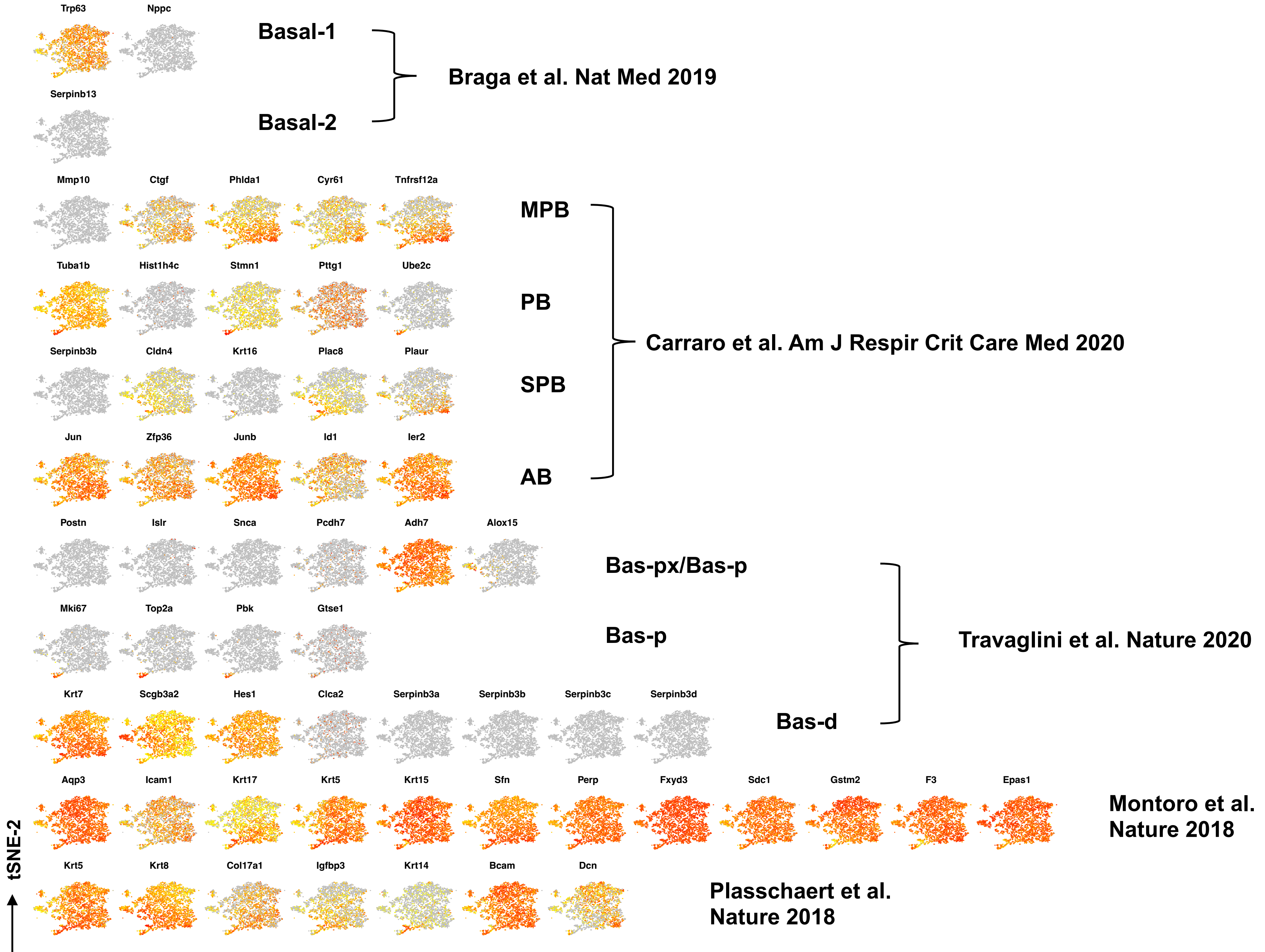
Trp63
Nppc
Serpinb13
Basal-1
Basal-2
Braga et al. Nat Med 2019
Mmp10
Ctgf
Phlda1
Cyr61
Tnfrsf12a
Tuba1b
Hist1h4c
Stmn1
Pttg1
Ube2c
Serpinb3b
Cldn4
Krt16
Plac8
Plaur
Jun
Zfp36
Junb
Id1
Ier2
MPB
PB
SPB
AB
Carraro et al. Am J Respir Crit Care Med 2020
Postn
Islr
Snca
Pcdh7
Adh7
Alox15
Mki67
Top2a
Pbk
Gtse1
Krt7
Scgb3a2
Hes1
Clca2
Serpinb3a
Serpinb3b
Serpinb3c
Serpinb3d
Bas-px/Bas-p
Bas-p
Bas-d
Travaglini et al. Nature 2020
Aqp3
Icam1
Krt17
Krt5
Krt15
Sfn
Perp
Fxyd3
Sdc1
Gstm2
F3
Epas1
Montoro et al.
Nature 2018
Krt5
Krt8
Col17a1
Igfbp3
Krt14
Bcam
Dcn
Plasschaert et al.
Nature 2018
tSNE-2
tSNE-1

Figure S3

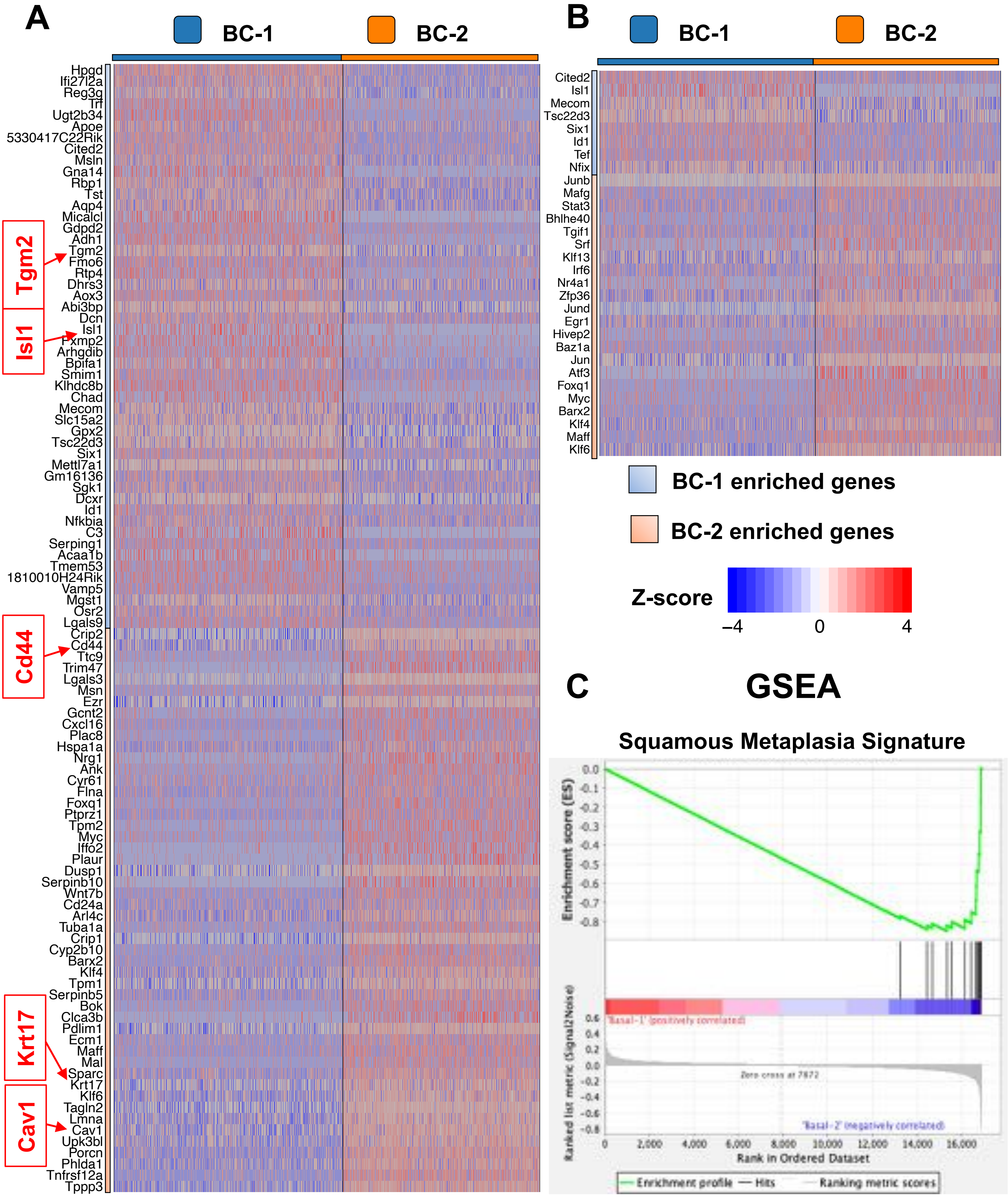
A
BC-1
BC-2
Hpgd
Ifi27l2a
Reg3g
Trf
Ugt2b34
Apoe
5330417C22Rik
Cited2
Msln
Gna14
Rbp1
Tst
Aqp4
Micalcl
Gdpd2
Adh1
Tgm2
Fmo6
Rtp4
Dhrs3
Aox3
Abi3bp
Dcn
Isl1
Fxmp2
Arhgdib
Bpifa1
Smim1
Klhdc8b
Chad
Mecom
Slc15a2
Gpx2
Tsc22d3
Six1
Mettl7a1
Gm16136
Sgk1
Dcxr
Id1
Nfkbia
C3
Serping1
Acaa1b
Tmem53
1810010H24Rik
Vamp5
Mgst1
Osr2
Lgals9
Crip2
Cd44
Ttc9
Trim47
Lgals3
Msn
Ezr
Gcnt2
Cxcl16
Plac8
Hspa1a
Nrg1
Ank
Cyr61
Flna
Foxq1
Ptprz1
Tpm2
Myc
Iffo2
Plaur
Dusp1
Serpinb10
Wnt7b
Cd24a
Arl4c
Tuba1a
Crip1
Cyp2b10
Barx2
Klf4
Tpm1
Serpinb5
Bok
Clca3b
Pdlim1
Ecm1
Maff
Mal
Sparc
Krt17
Klf6
Tagln2
Lmna
Cav1
Upk3bl
Porcn
Phlda1
Tnfrsf12a
Tppp3
Tgm2
Isl1
Cd44
Krt17
Cav1
B
BC-1
BC-2
Cited2
Isl1
Mecom
Tsc22d3
Six1
Id1
Tef
Nfix
Junb
Mafg
Stat3
Bhlhe40
Tgif1
Srf
Klf13
Irf6
Nr4a1
Zfp36
Jund
Egr1
Hivep2
Baz1a
Jun
Atf3
Foxq1
Myc
Barx2
Klf4
Maff
Klf6
BC-1 enriched genes
BC-2 enriched genes
Z-score
–4
0
4
C
GSEA
Squamous Metaplasia Signature
Enrichment score (ES)
0.0
-0.1
-0.2
-0.3
-0.4
-0.5
-0.6
-0.7
-0.8
'Basal-1' (positively correlated)
Zero cross at 7872
'Basal-2' (negatively correlated)
Ranked list metric (Signal2Noise)
0.6
0.4
0.2
0.0
-0.2
-0.4
-0.6
-0.8
0
2,000
4,000
6,000
8,000
10,000
12,000
14,000
16,000
Rank in Ordered Dataset
Enrichment profile
Hits
Ranking metric scores

**Figure S4**

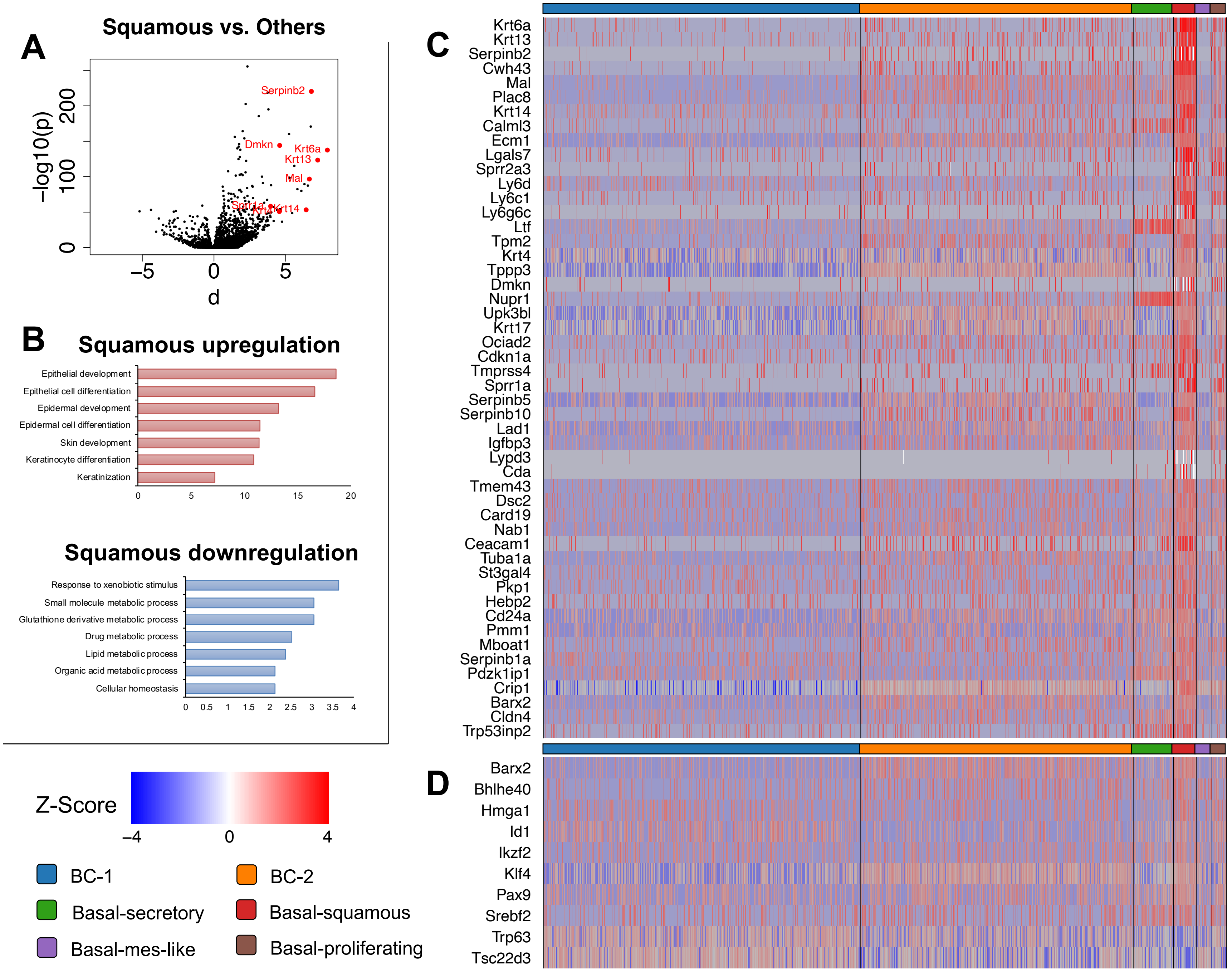

A
Squamous vs. Others
-log10(p)
d
Serpinb2
Dmkn
Krt6a
Krt13
Mal
Sprr1a
Krt4
Krt14
B
Squamous upregulation
Epithelial development
Epithelial cell differentiation
Epidermal development
Epidermal cell differentiation
Skin development
Keratinocyte differentiation
Keratinization
Squamous downregulation
Response to xenobiotic stimulus
Small molecule metabolic process
Glutathione derivative metabolic process
Drug metabolic process
Lipid metabolic process
Organic acid metabolic process
Cellular homeostasis
Z-Score
BC-1
BC-2
Basal-secretory
Basal-squamous
Basal-mes-like
Basal-proliferating
C
Krt6a
Krt13
Serpinb2
Cwh43
Mal
Plac8
Krt14
Calml3
Ecm1
Lgals7
Sprr2a3
Ly6d
Ly6c1
Ly6g6c
Ltf
Tpm2
Krt4
Tppp3
Dmkn
Nupr1
Upk3bl
Krt17
Ociad2
Cdkn1a
Tmprss4
Sprr1a
Serpinb5
Serpinb10
Lad1
Igfbp3
Lypd3
Cda
Tmem43
Dsc2
Card19
Nab1
Ceacam1
Tuba1a
St3gal4
Pkp1
Hebp2
Cd24a
Pmm1
Mboat1
Serpinb1a
Pdzk1ip1
Crip1
Barx2
Cldn4
Trp53inp2
D
Barx2
Bhlhe40
Hmga1
Id1
Ikzf2
Klf4
Pax9
Srebf2
Trp63
Tsc22d3


**Figure S5**

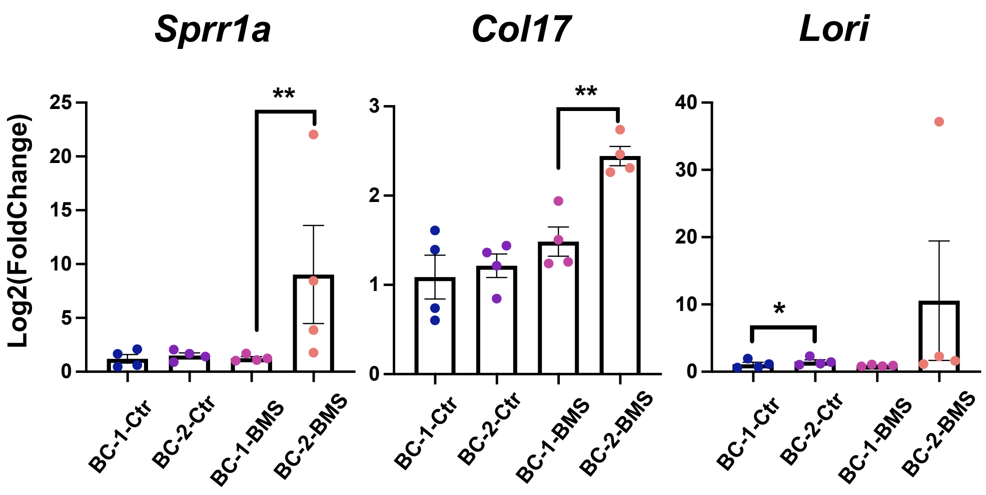

Sprr1a
Col17
Lori
Log2(FoldChange)
25
20
15
10
5
0
3
2
1
0
40
30
20
10
0
**
**
*
BC-1-Ctr
BC-2-Ctr
BC-1-BMS
BC-2-BMS


**Figure S6**

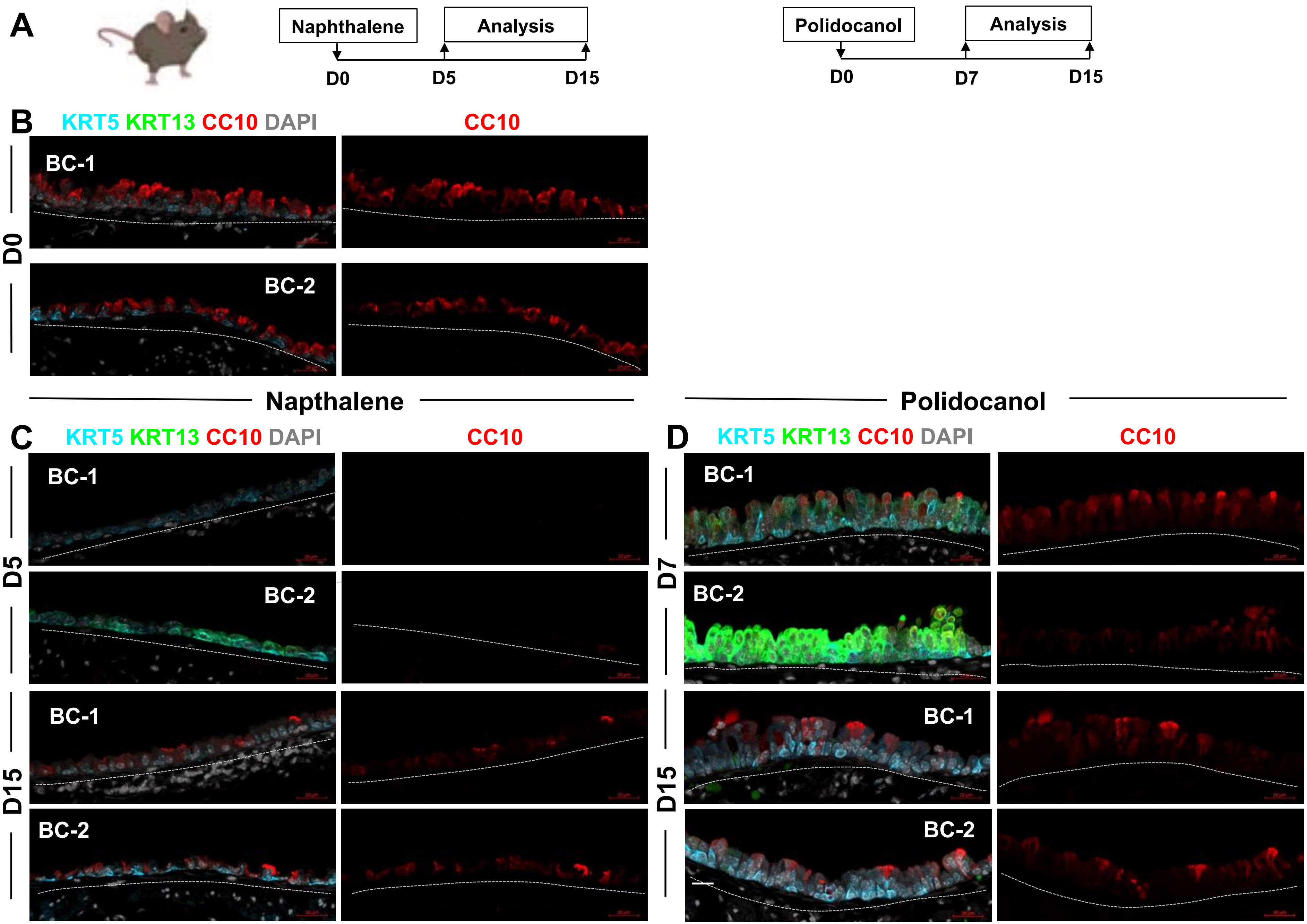
A
Naphthalene
Analysis
D0
D5
D15
Polidocanol
Analysis
D0
D7
D15
B
KRT5 KRT13 CC10 DAPI
CC10
BC-1
BC-2
D0
Napthalene
Polidocanol
C
KRT5 KRT13 CC10 DAPI
CC10
BC-1
BC-2
BC-1
BC-2
D5
D15
D
KRT5 KRT13 CC10 DAPI
CC10
BC-1
BC-2
BC-1
BC-2
D7
D15

Figure S7
